# The effects of increasing the number of taxa on inferences of molecular convergence

**DOI:** 10.1101/081612

**Authors:** Gregg W.C. Thomas, Matthew W. Hahn, Yoonsoo Hahn

## Abstract

Convergent evolution provides insight into the link between phenotype and genotype. Recently, large-scale comparative studies of convergent evolution have become possible, but researchers are still trying to determine the best way to design these types of analyses. One aspect of molecular convergence studies that has not yet been investigated is how taxonomic sample size affects inferences of molecular convergence. Here we show that increased sample size decreases the amount of inferred molecular convergence associated with the three convergent transitions to a marine environment in mammals. The sampling of more taxa—both with and without the convergent phenotype—reveals that alleles associated only with marine mammals in small datasets are actually more widespread, or are not shared by all marine species. The sampling of more taxa also allows finer resolution of ancestral substitutions, revealing that they are not in fact on lineages leading to solely marine species. We revisit a previous study on marine mammals and find that only 7 of the reported 43 genes with convergent substitutions still show signs of convergence with a larger number of background species. However, 4 of those 7 genes also showed signs of positive selection in the original analysis and may still be good candidates for adaptive convergence. Though our study is framed around the convergence of marine mammals, we expect our conclusions on taxonomic sampling are generalizable to any study of molecular convergence.

## Introduction

From the beginning of the genome era, researchers have attempted to associate individual genomic features with the phenotypes present in sequenced species. Such features include unique amino acid substitutions (Kim et al. 2011), gene family expansions (Lespinet et al. 2002) or contractions (Aravind et al. 2000), and changes in gene expression (Khaitovich et al. 2005), to name a few. In order to increase the power to detect causative changes as more genomes were sequenced, researchers looked for convergent molecular changes associated with convergent phenotypes. Again, such changes were found among amino acid substitutions (e.g. Rokas and Carroll 2008; Foote et al. 2015), gene gains and losses (e.g. Pellegrini et al. 1999; McBride and Arguello 2007; Hiller et al. 2012; De Smet et al. 2013; Liebeskind et al. 2015), in sex chromosome morphology (e.g. Bellott et al. 2010), and in gene expression (e.g. Ogura et al. 2004; Pankey et al. 2014).

However, it has become evident that the number of species and genomes used in an analysis greatly influences the probability of finding unique genomic changes in only taxa with the phenotypes of interest. For example, the original description of the naked mole-rat genome found that a unique amino acid substitution may have been responsible for the rodent’s hairless phenotype (Kim et al. 2011). A subsequent study with a larger number of taxa found that this substitution was not, in fact, unique to naked mole-rats but is found throughout mammals, including in the closest living relative to the naked mole-rat, the guinea pig (Delsuc and Tilak 2015). Similarly, we expect that as more genomes are analyzed in studies looking for uniquely convergent molecular changes—those shared only by the taxa with convergent phenotypes—it will be more and more likely that we find similar changes in non-phenotypically convergent species. While such analyses will certainly increase our power to identify truly causative convergent mutations, in some cases they may also reveal that convergent phenotypes are not underlain by convergent molecular mechanisms (Storz 2016).

Here we investigate how the number of taxa sampled affects inferences of convergent evolution among amino acid sequences. We find that as more lineages are added to an analysis, the number of unique convergent substitutions among a given set of species decreases rapidly. We also find that adding taxa can decrease the number of convergent substitutions along target lineages (unique to those lineages or not), as the added phylogenetic resolution can help to more accurately reconstruct ancestral sequences. Finally, we revisit results on convergent amino acid substitutions among marine mammals presented in Foote et al. (2015) to see how these substitutions hold up to the addition of more species to the analysis. Three separate mammalian lineages have transitioned to an aquatic lifestyle and in the process have experienced extensive phenotypic convergence (Kelley et al. 2016), making them a prime choice for the study of convergence at the molecular level.

## Methods

To assess how the number of taxa influences inferences of molecular convergence, we began by downloading the 100 species hg19 human reference alignments from the UCSC Genome Browser (Kent et al. 2002; http://hgdownload.soe.ucsc.edu/goldenPath/hg19/multiz100way/). We pruned and filtered all but 59 mammal species (Figure 1). Of these 59 species, five are marine mammals. The West Indian manatee (*Trichechus manatus latirostris*) is from the Order Sirenia; two Cetacean species are present, the killer whale (*Orcinus orca*) and the bottlenose dolphin (*Tursiops truncatus*); and there are two Pinniped species, the walrus (*Odobenus rosmarus*) and the Weddell seal (*Leptonychotes weddellii*). These three orders represent three independent gains of phenotypes associated with aquatic lifestyles in mammals (Figure 1; Foote et al. 2015), and were the target lineages for our tests of convergence. In our analyses, we classify lineages in which we are searching for convergence as “target” lineages and all others within the phylogeny as “background” lineages. We filtered all but the longest isoform from the original 39,361 sequences giving us 19,215 orthologs from which to draw on for our analysis of unique convergent substitutions.

**Figure 1:**
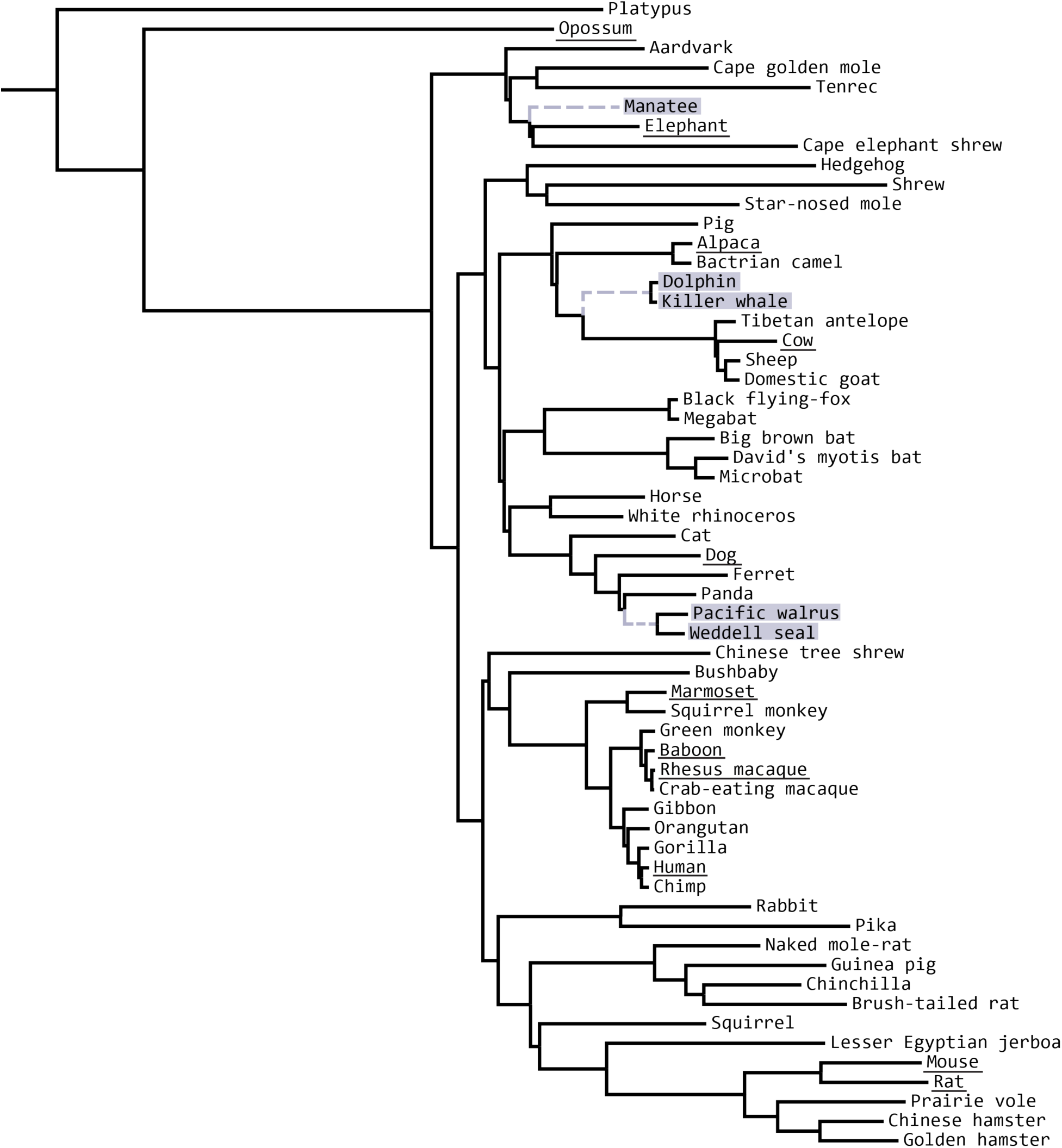
The 59 mammal species used in the analyses. Marine mammals are highlighted in grey and were used as target species to count unique convergent substitutions. Branches leading to marine mammal clades are dotted and grey and were used as target lineages to count convergent substitutions with ancestral sequence reconstruction. The 11 terrestrial mammal species used in Foote et al. (2015) are underlined.

We define unique convergent substitutions as sites where all marine mammal species have an identical amino acid that is different from any amino acid found in the background species. We count unique convergent substitutions with varying numbers of background species in the following way: starting with the five marine mammal species, we randomly added *n* background species and counted unique convergent sites for each dataset. Values of *n* range from 11 to 54 (there were 11 background species in the original analysis of Foote et al. 2015), and we repeated the process 100 times for each *n*.

We also identified possible convergent substitutions by reconstructing ancestral amino acid states and looking for changes along the lineages leading to the three marine mammal clades (grey, dashed branches in Figure 1). For this analysis we restricted our set of 19,215 sequences to only those 10,521 in which all 59 mammalian species were present. We used *codeml* in PAML (Yang 2007) to reconstruct ancestral sequences and counted a site as a convergent substitution if a change from any ancestral state to the descendant state had occurred along all three of these branches and resulted in the same descendant state. We followed a similar replicate scheme as above, but with fewer values of *n* and fewer numbers of replicates due to computational constraints. Values of *n* for ancestral reconstructions are: 15, 20, 25, 30, 35, 40, 45, and 50 and the process was repeated 5 times for each value of *n*. For this experiment, background species are chosen randomly for every replicate at every value of *n*.

Finally, we replicated the analysis in Foote et al. (2015) by performing ancestral reconstructions on the same species used in that study (highlighted and underlined species in Figure 1, excluding Weddell seal) and counting convergent substitutions along the marine mammal lineages. We then added Weddell seal to the analysis to see how another target species affects inferences of convergence. From there, we iteratively added background species (15, 20, 25, 30, 35, 40, 45, 50 species) and repeated the convergence inference process. In this iterative setup, the background species that are added are still chosen randomly, but all species in previous replicates are included in the current one. For example, in the replicate with 20 background species, 15 of them are preserved from the trial with 15 background species and 5 more are added randomly. At each replicate, the list of genes with convergent substitutions was compared to the list of 43 genes with convergent substitutions inferred by Foote et al. (2015).

## Results

Our current study is motivated by the recent investigation of molecular convergence in marine mammals by Foote et al. (2015). In that study, the authors (including G.W.C.T. and M.W.H.) used the genomes of four marine mammals (highlighted in Figure 1, excluding the Weddell seal) as target species and 11 terrestrial mammals (underlined in Figure 1) as background species. Alignments of those 15 species were used to identify convergent substitutions by inferring ancestral sequences. In total, convergent substitutions in 43 genes were found along the lineages leading to all three marine mammal clades. We used the same number of species as the starting point for this study.

### Sample size affects the number of uniquely convergent substitutions

We find that as the number of background taxa is increased, the number of uniquely convergent substitutions among a set of target species decreases (these represent amino acid states found only in marine mammals). We observed this by counting unique convergent substitutions among the five marine mammal species highlighted in Figure 1 and randomly adding other background mammalian taxa to the analysis in increasing numbers (Figure 2). Although there is a lot of variability in the exact number of initially unique convergent substitutions that are found in other lineages—and this is associated with the random sampling of additional background lineages—there is a clear monotonic relationship that appears to approach zero. That is, our results imply that with enough additional taxa added there will be no unique convergent substitutions among the three marine mammal clades. Using all 54 background species, we find only 1 uniquely convergent substitution in one gene (*ITGA8*; NM_003638). (Figure 2).

**Figure 2:**
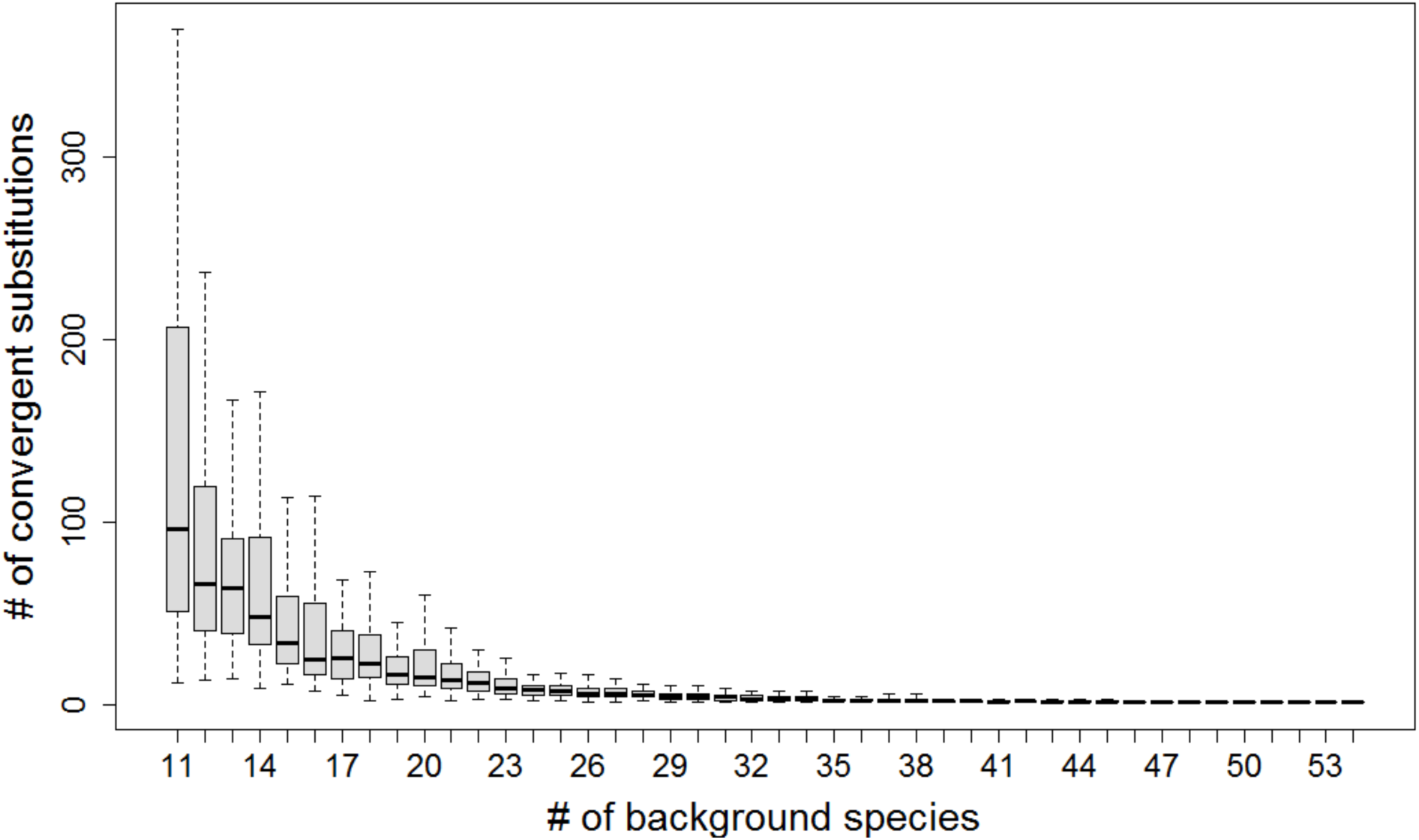
The number of unique convergent substitutions counted among marine mammal species rapidly decreases with an increasing number of background species.

We also tested how the number of target species affects counts of unique substitutions by specifying the same target and background species used in Foote et al. (2015) and either adding Weddell seal or removing killer whale or dolphin. With the original 15 species, we find 131 unique convergent substitutions. However, when Weddell seal is added as a target species this number drops to 87. This is because the seal has a different amino acid at a position where the other four marine mammals share one to the exclusion of all other species in the analysis. Likewise, when either killer whale or dolphin are removed from the analysis as target species, the number of uniquely convergent substitutions rises to 144 and 141, respectively. In this case the removal of target species has made it more likely to find amino acids shared among a smaller set of marine mammals.

### Sample size affects inferences of convergent evolution by changing ancestral reconstructions

Molecular convergence can also be assessed by using ancestral sequence reconstruction to look for shared changes along specific branches of interest in a phylogeny. Following this procedure, we searched for convergent substitutions along the three branches leading to marine mammals in our dataset (grey, dashed branches in Figure 1) while varying the number of background species. We again find a sharp drop-off in convergent substitutions with increasing numbers of background species (Figure 3). Note that unlike in the previous analysis, here it is the reconstruction that has changed, not the uniqueness of these convergent substitutions. This analysis solely asks whether the substitutions are convergent along all three lineages based on ancestral sequence reconstruction, not whether additional substitutions to the same state have occurred elsewhere on the tree.

**Figure 3:**
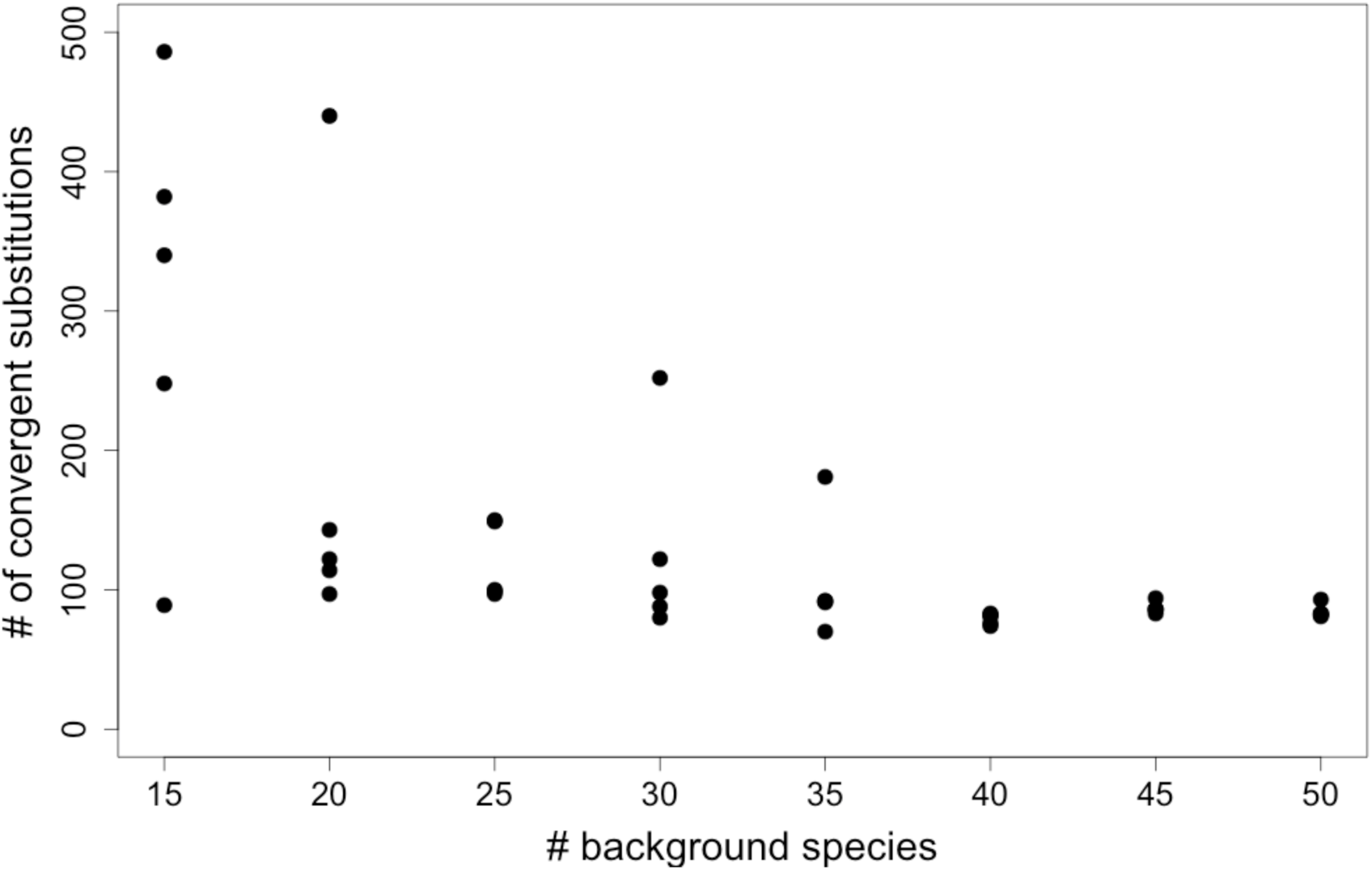
The number of convergent substitutions along marine mammal lineages as inferred through ancestral sequence reconstruction rapidly decreases with an increasing number of background species.

The apparent loss of convergent substitutions is caused when the inclusion of additional species changes the reconstructed ancestral amino acid states, leading the substitutions to be assigned to different branches. In the many cases we examined this occurs because the reconstructed state at the ancestral nodes of the target lineages is inferred to have the state shared among marine mammals. Therefore, no convergent substitution along these three branches is necessary. Figure 4 highlights an example in which a convergent substitution is found in marine mammals in an analysis with 11 background species (serine → lysine or alanine → lysine). However, with the addition of four more species, three of which have an observed lysine residue at that position, the ancestral reconstructions throughout the tree are changed, mostly to lysine to accommodate the new observed states. The original convergent substitution is therefore no longer counted, as it has been moved to a non-marine mammal lineage.

**Figure 4:**
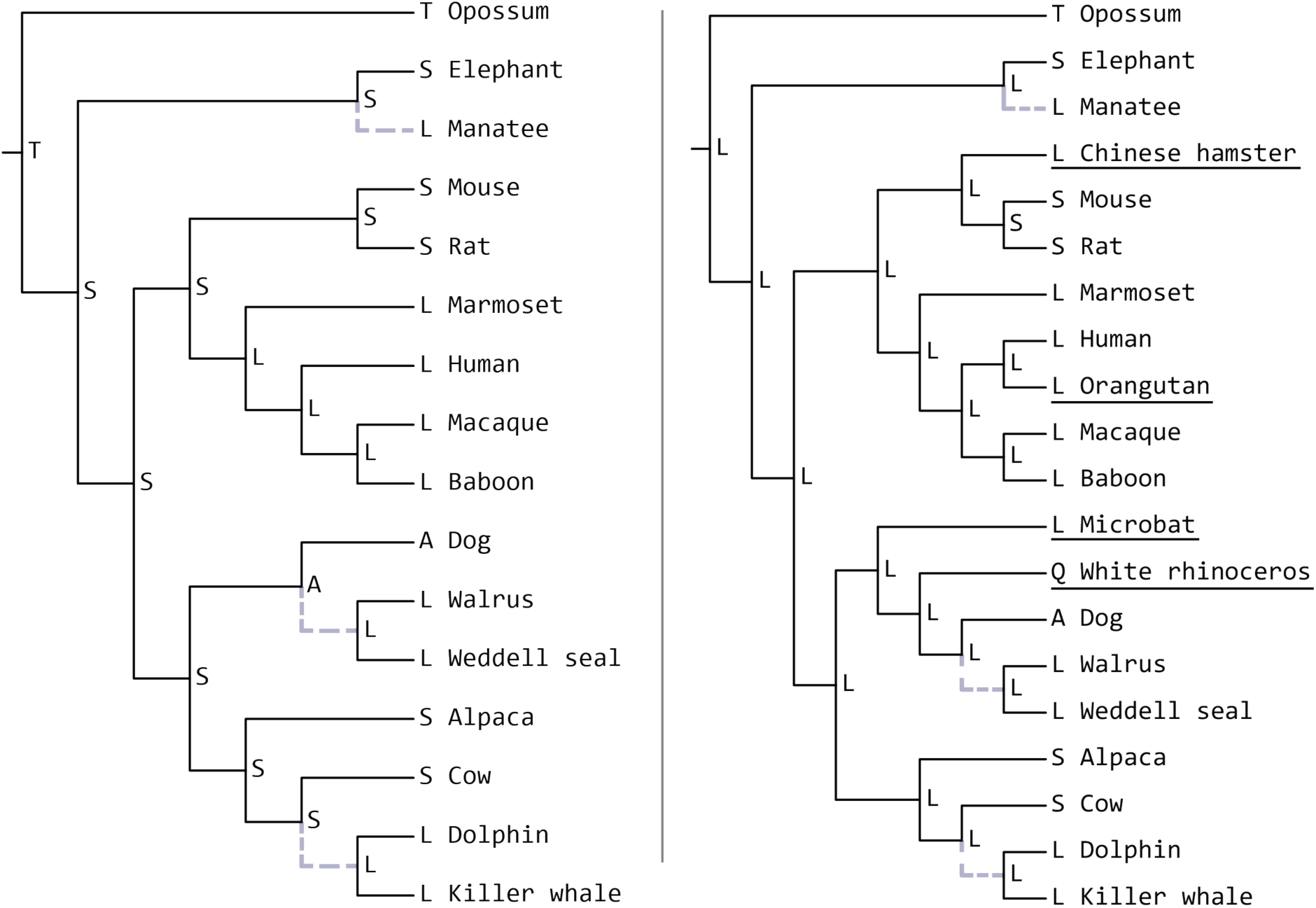
An example of loss of the signal of convergence for a single gene, NM_000062. (***Left***) With these 16 species the three marine mammal lineages (grey and dashed branches) all have convergent substitutions from some ancestral state to lysine (L). (***Right***) The addition of four more species (underlined) changes ancestral states throughout the tree. Now states towards the root appear as lysine (L) rather than serine (S), meaning changes to lysine (L) are no longer needed along the branches leading to the marine mammals.

### Comparisons with the original analysis of marine mammal convergence

Given these results, we revisited the original findings of amino acid convergence among marine mammals (Foote et al. 2015). In that paper, the authors found 44 convergent substitutions in 43 genes along marine mammal lineages among a set of 5,893 orthologs. Replicating their analysis (using ancestral sequence reconstruction and the same set of species) with our set of 10,521 genes, we find 246 convergent substitutions in 233 genes. 42 of the original 43 genes with convergent substitutions are present in our dataset, yet we were surprised to find only 27 with convergent substitutions in our analysis. This indicates that the alignment procedure may also affect inferences of convergent substitutions. The pattern of decreased convergence shared by the marine mammals continues as we add additional background species to the analysis: the total number of convergent substitutions and the number of convergent genes recovered from the original study decreases (Table 1).

**Table 1:**
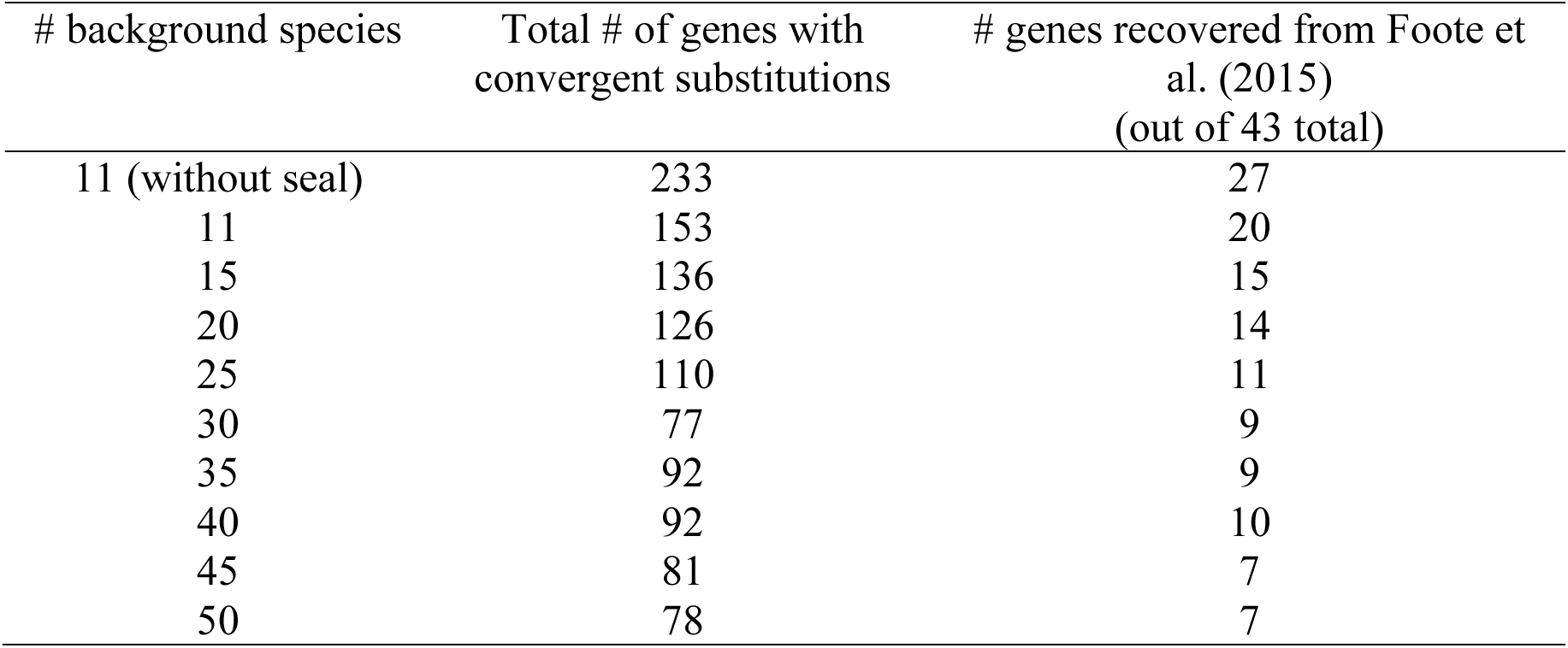
The number of convergent substitutions as inferred by ancestral sequence reconstruction in marine mammals with increasing numbers of background species.

| # background species | Total # of genes with convergent substitutions | # genes recovered from Foote et al. (2015)<br>(out of 43 total) |
| --- | --- | --- |
| 11 (without seal) | 233 | 27 |
| 11 | 153 | 20 |
| 15 | 136 | 15 |
| 20 | 126 | 14 |
| 25 | 110 | 11 |
| 30 | 77 | 9 |
| 35 | 92 | 9 |
| 40 | 92 | 10 |
| 45 | 81 | 7 |
| 50 | 78 | 7 |

When we add the fifth marine mammal, Weddell seal, to the analysis, the number of convergent substitutions drops to 161 and we recover only 20 of the original 43 convergent genes found by Foote et al. (2015). The decrease in convergent substitutions, even putatively adaptive ones, with more focal taxa is not unexpected. Foote et al. (2015) previously reported that some convergent substitutions in the four marine mammals studied were not present in further comparisons with the minke whale genome (Yim et al. 2013), some were not. However, because minke was not included in their phylogenetic analysis, it is possible that these substitutions represent reversions in minke whale.

We also replicated Foote et al.’s (2015) use of terrestrial mammals as an empirical null distribution to test if the amount of convergence in marine mammals exceeds expectations. In the original paper, the authors counted convergent substitutions between dog, cow, and elephant as a control and found no excess of convergence in marine mammals compared to these land mammals. The authors in fact reported more convergence in land mammals, with 93 convergent substitutions in 90 genes. Similarly, we also find that convergence in marine mammals does not exceed that of convergence in the same set of land mammals, even when increasing the number of taxa in the dataset. With the original set of species we find convergent substitutions in 398 genes among land mammals, many more than the 233 in marine mammals. This pattern holds with increasing numbers of background species. With all 50 background species, we find 117 genes with convergent substitutions in land mammals compared to 78 in marine mammals (Table 1). This confirms the original findings of Foote et al. (2015) that there is no excess amino acid convergence among marine mammals. This also implies that the pattern of decreasing convergence with increasing number of taxa remains regardless of the specific target species used in the phylogeny.

Interestingly, several of the genes originally found as convergent by Foote et al. (2015) continue to show signals of convergence despite the increased number of background species (Table 2). Some of these genes also show signs of positive selection in the original analysis (based on analyses of *d*_N_/*d*_S_) and have possible functional implications for phenotypic convergence among marine mammals. *MGP9* may play a role in bone formation, *MYH7B* is involved in cardiac muscle development, and *SERPINC1* regulates blood coagulation. *GCLC* is another interesting gene that we still detect as convergent with a large number of species. This gene is involved in glutathione metabolism, a molecule that has been shown to prevent oxidative damage in cetaceans during long underwater dives.

**Table 2:** Genes found to be convergent by Foote et al. (2015) and when using 50 background species in this study.

| Gene name | UCSC Gene ID | Evidence for positive selection<br>in Foote et al. (2015) |
| --- | --- | --- |
| *ZNF292* | NM_015021 |  |
| *MYH7B* | NM_020884 | * |
| *GCLC* | NM_001197115 | * |
| *SERPINC1* | NM_000488 | * |
| *DUSP27* | NM_001080426 |  |
| *M6PR* | NM_002355 | * |
| *SIAE* | NM_170601 |  |

## Discussion

The study of convergence at the molecular level is still relatively new. Although most recent papers have examined convergence in amino acid substitutions (e.g. Rokas and Carroll 2008; Foote et al. 2015), there can be many types of convergent molecular changes, including gene gains, losses, and shifts in gene expression (e.g. De Smet et al. 2013; Liebeskind et al. 2015; Ogura et al. 2004; Pankey et al. 2014). Researchers are also discovering (or re-discovering) many nuances that must be considered when measuring molecular convergence, such as choosing the correct null model (Thomas and Hahn 2015; Zou and Zhang 2015a) and accounting for physical constraints on convergence (Goldstein et al. 2015; Zou and Zhang 2015b). The number of taxa in an experiment can also affect any possible inferences of lineage-specific changes (Delsuc and Tilak 2015) and this must also be considered in measures of molecular convergence.

Stayton (2008) showed that inferences of convergence in quantitative traits increased with increasing number of taxa, and this must also be true of molecular data. With the addition of each new species, there will also be more branches in the phylogeny on which substitutions of all types can occur. What we have shown here is that increased sample size decreases inferences of molecular convergence *among a set of target species*. These results illustrate the need for caution when making conclusions about molecular convergence. Molecular convergence that is responsible for adaptive phenotypic convergence appears to be much less common when additional species are included in any analysis. In many cases, the convergent amino acid state will be sampled in a non-phenotypically convergent species. In our study of marine mammals, these species were not even semi-aquatic mammals, thus ruling out these changes as being adaptively convergent (even if they are). Similar patterns of ecologically diverse taxa sharing convergent states are found in studies of morphological characters (Zou and Zhang 2016).

We expect to see the same effect of taxonomic sampling when considering convergence in any type of molecular change. As more species are added, for instance, more gene duplications and losses will have occurred, making it likely that we observe such changes in non-phenotypically convergent lineages. There can also be different ways that changes in protein sequences can be convergent. Chikina et al. (2016) propose that molecular convergence may be the result of selection at the gene level rather than the amino acid level. These authors found an excess of genes with accelerated or decelerated rates of evolution across marine mammals when compared to terrestrial mammals. This model proposes that multiple different changes in common genes lead to a convergent phenotype rather than a single identical convergent amino acid change. We expect that this measure of convergence should still be sensitive to changes in the number of species represented, though we have not tested this here (note that Chikina et al. use at least 30 species in their analysis).

In this study, we used two methods for detecting convergent substitutions: unique substitutions among a set of target species and convergent changes inferred with ancestral reconstructions on a set of target lineages. Both methods showed a similar pattern of decreasing numbers of convergent sites with increasing taxa, though they may be affected by different features of the added taxa. The use of unique substitutions would seem to be a conservative method, as the presence of “convergent” amino acid states in non-phenotypically convergent species does not mean such changes are not involved in adaptation. For example, if the individual sequenced from a newly added species carries a deleterious allele at mutation-selection balance at a site that is truly convergent in a set of target species, we would eliminate this substitution as a candidate. Convergent substitutions inferred with ancestral reconstructions may be lost because additional taxa move the convergent substitution to a different branch of the phylogeny, or because they reverse the direction of the evolutionary change (e.g. Figure 4).

Ancestral reconstructions are also dependent on the tree topology. The addition of new taxa may change the species tree topology and drastically alter inferred ancestral states. Though we have not explored the effect of additional taxa on improving the underlying topology, this will clearly affect inferences of convergence. Incorrect topologies—either due to error in species tree inference or biological differences in the histories among different genes—can lead to truly convergent substitutions being missed, but can also lead to many incorrect inferences of convergence (Mendes and Hahn 2016; Mendes, Hahn, and Hahn *in press*).

Aside from the quantification of molecular convergence, an important consideration in any study is quantifying whether there is an excess amount of convergence. Because molecular convergence can be the result of neutral processes (Zhang and Kumar 1997), appropriate null comparisons must be made. The best way to account for this background convergence is with the use of an empirical null model, such as comparisons among non-convergent species that are closely related to the target lineages. For example, Foote et al. (2015) found no evidence of excess convergence in marine mammals given levels of background convergence in the terrestrial species elephant, dog, and cow. We find similar results here with our larger taxonomic sample. And while the signal for convergence remained in several of the interesting genes originally found by Foote et al. (Table 2), we must point out that even with 50 species the trend does not seem to have levelled out. This could indicate that perhaps with even more species some of these genes would no longer be convergent.

We recommend that future studies of molecular convergence should be performed so that they maximize the number of taxa represented for more confident inferences. Our results show that adding either target or background species can change the outcome of convergence analyses, so it is currently difficult to determine which is more valuable. However, it is likely that adding background species will be easier in most cases since target species sharing a convergent trait will be less common.

## Acknowledgments

We would like to thank Andy Foote for helpful feedback with the manuscript. We acknowledge computational resources provided by the National Center for Genome Analysis Support (National Science Foundation grant DBI-1458641).

